# The giant staphylococcal protein Embp facilitates colonization of surfaces through Velcro-like attachment to fibrillated fibronectin

**DOI:** 10.1101/2021.05.31.446437

**Authors:** Nasar Khan, Hüsnü Aslan, Henning Büttner, Holger Rohde, Thaddeus Wayne Golbek, Steven Joop Roeters, Sander Woutersen, Tobias Weidner, Rikke Louise Meyer

**Affiliations:** Interdisciplinary Nanoscience Center (iNANO), Aarhus University, 8000 Aarhus, Denmark; Institute for Medical Microbiology, Virology and Hygiene, University Medical Centre Hamburg-Eppendorf, 20246 Hamburg, Germany; Department of Chemistry, Aarhus University, 8000 Aarhus C, Denmark; Van ‘t Hoff Institute of Molecular Sciences, University of Amsterdam, 1098XH, Amsterdam, the Netherlands; Department of Biology, Aarhus University, 8000 Aarhus C, Denmark

## Abstract

*Staphylococcus epidermidis* causes some of the most hard-to-treat clinical infections by forming biofilms: Multicellular communities of bacteria encased in a protective matrix, supporting immune evasion and tolerance against antibiotics. Biofilms occur most commonly on medical implants, and a key event in implant colonization is the robust adherence to the surface, facilitated by interactions between bacterial surface proteins and host matrix components. *S. epidermidis* is equipped with a giant adhesive protein, Embp, which facilitates bacterial interactions with surface-deposited, but not soluble fibronectin. The structural basis behind this selective binding process has remained obscure. Using a suite of single-cell and single-molecule analysis techniques, we show that *S. epidermidis* is capable of such distinction because Embp binds specifically to fibrillated fibronectin on surfaces, while ignoring globular fibronectin in solution. *S. epidermidis* adherence is critically dependent on multi-valent interactions involving 50 fibronectin-binding repeats of Embp. This unusual, Velcro-like interaction proved critical for colonization of surfaces under high flow, making this newly identified attachment mechanism particularly relevant for colonization of intravascular devices, such as prosthetic heart valves or vascular grafts. Other biofilm-forming pathogens, such as *Staphylococcus aureus*, express homologs of Embp and likely deploy the same mechanism for surface colonization. Our results may open for a novel direction in efforts to combat devastating, biofilm-associated infections, as the development of implant materials that steer the conformation of adsorbed proteins is a much more manageable task than avoiding protein adsorption altogether.

**Graphical abstract:** Fibronectin exists in two different conformations in the body. It circulates in the bodily fluids in globular conformation, however, it become fibrillated once adsorbed to an implant surface. *S. epidermidis* possess a giant 1 MDa receptor known as Embp bind specifically to fibrillated Fn but not to the globular Fn.

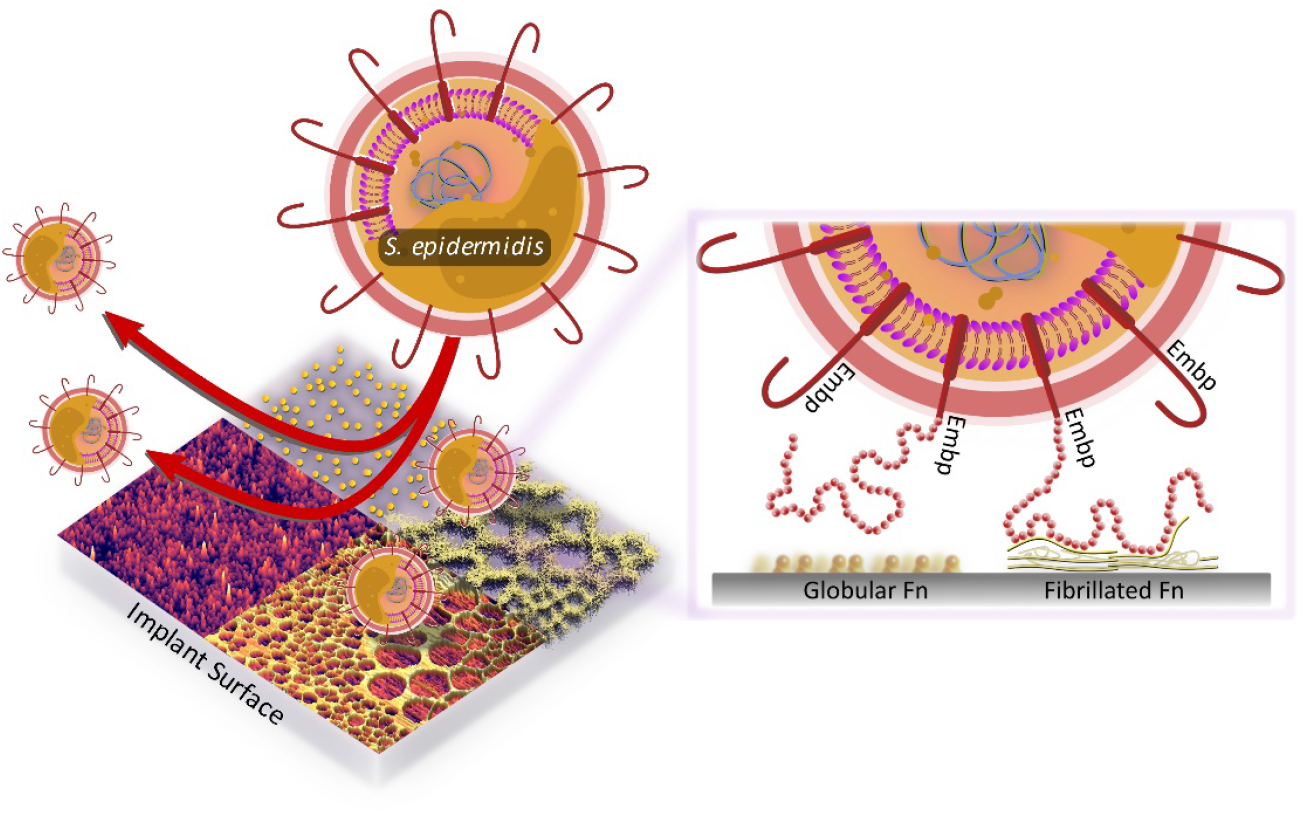

## INTRODUCTION

Biomedical implants, such as catheters, prosthetics, vascular grafts, and similar devices have revolutionized the medical field. However, implants can lead to severe infections due to bacterial biofilms: The formation of multicellular bacterial communities encased in a protective extracellular matrix (*1*). Bacteria in the biofilm evade phagocytosis by immune cells (*2*), and the immune system can therefore not eradicate the infection. Furthermore, a fraction of the cells enter a dormant state in which they are highly tolerant to antibiotics (*3*). With the rise in use of biomedical implants, there is an urgent and growing need to understand how biofilm infections arise, such that new strategies for preventative treatment can be developed.

Staphylococci, particularly *Staphylococcus aureus* and *Staphylococcus epidermidis* are the culprits of most implant-associated infections (*4*). Despite its low virulence, *S. epidermidis* is common in these infections due to its prowess in biofilm formation. *S. epidermidis* attaches to implant surfaces via adsorbed host proteins (*5, 6*) and it expresses an array of surface-bound proteins (adhesins) that bind to host proteins, such as fibrin, fibronectin, vitronectin, and collagen to initiate biofilm formation (*7*). One such adhesin is the extracellular matrix binding protein (Embp), which is found in the vast majority of clinical isolates of *S. epidermidis* (*8, 9*), suggesting that this giant 1 MDa protein is important for this species’ pathogenicity. Embp contains a number of repetitive motifs. These were originally described based on sequence similarity to be 21 “Found in Various Architecture” (FIVAR) repeats and 38 alternating repeats of “G related Albumin Binding” (GA) and FIVAR repeats combined, named FIVAR-GA repeats. After the crystal structure was recently solved, the domain structure was updated and consists of 10 170-aa F-repeats that each represent two FIVAR repeats, and 40 125-aa FG-repeats that each represent the previously termed FIVAR-GA repeats (*10*). These 50 repeats can bind to fibronectin (Fn), and it is presumed that this interaction aids the colonization of the host (*11*).

The Fn deposition can occur around implants (*12*) and offers a site for bacterial attachment. We wondered how bacteria like *S. epidermidis* can colonize implant surfaces by interacting with adsorbed Fn when the same protein is also abundant in a soluble form in blood. Presumably, Fn-binding proteins on the bacterial cell surface become occupied with soluble Fn before being able to interact with Fn on the implant surface. The aim of this study was to determine how pathogens overcome this dilemma and bind to host proteins on tissue or implant surfaces while ignoring soluble forms of the same protein. Understanding the pathogens’ ability to selectively colonize implant surfaces reveals conceptual mechanisms for how pathogens control their location and fate in the host.

In this study, we investigate Embp’s interaction with Fn. Fn circulates in bodily fluids in a compact globular form (*13*), while fibrillated Fn contributes to the assembly of the extracellular matrix of tissue (*14*). It is the stretching of Fn upon interacting with cell surface integrins, which exposes self-binding domains and trigger Fn fibrillation. This mechanism ensures that Fn only fibrillates in the extracellular matrix of tissue and not in the blood stream (*15*). We hypothesize that *S. epidermidis* interacts selectively with fibrillated Fn, and that a fibrillated ligand provides an opportunity for a multi-valent interaction with the many repetitive F and FG repeats of the Embp. Using a model system of polymer-coated surfaces that facilitate Fn adsorption in either globular or fibrillated conformation, we probed Embp’s interaction with Fn. Using native and recombinant Embp in a series of analyses at the population-, single-cell-, and single-molecule levels, we confirmed that Embp selectively interacts with fibrillated Fn. The interaction is a Velcro-like mechanism where multiple binding-domains must interact simultaneously to facilitate strong attachment. Such strong attachment via a single protein is particularly beneficial under high sheer stress, such as in the vascular system, and it was exactly under these conditions that Embp gave the cells and advantage. Embp homologs are present in other important pathogens capable of biofilm formation in the vascular system, and our study reveals a mechanism for how bacteria accomplish this feat.

## RESULTS

### Embp does not interact with soluble fibronectin

We hypothesized that Embp selectively binds to fibrillated Fn, which would allow the bacteria to colonize surfaces via Fn without being blocked by soluble Fn in the bloodstream. To study the interaction between Embp and Fn, we expressed Embp fusion proteins comprised of either 5 F-repeats (Embp_5F) or 9 FG-repeats (Embp_9FG), each fused to the native export signal and anticipated C-terminal cell wall anchor region (*10*), in the surrogate host *Staphylococcus carnosus* TM300, which has no other mechanisms for attachment to Fn. The full-length Embp is too large to clone into a surrogate host, and it was therefore not possible to investigate the full-length Embp protein. However, expression of the two different Embp fragments allowed us to study their interactions individually. The presence of these fragments on the cell surface was confirmed by immunofluorescence staining (Figure S1).

Neither F-nor FG-repeats facilitated adsorption of soluble fluorescently conjugated Fn to the surface of *S. carnosus* (Figure S2). The native Embp expressed by *S. epidermidis* did not bind soluble Fn either (Figure S2), concluding that Embp does not interact with Fn in its soluble, globular conformation.

### Embp interacts exclusively with fibrillated fibronectin

In order to further investigate Embp’s interaction with Fn in different conformations, we produced a model system in which Fn was adsorbed to a surface in either the globular of fibrillated conformation. Previous research had shown that Fn fibrillates when adsorbed on surfaces coated with poly (ethyl acrylate) (PEA), while it remains globular on poly (methyl acrylate) (PMA) (*16-18*). The two polymer coatings have similar physico-chemical properties, but the ethyl side group of PEA provides sufficient mobility of the adsorbed protein to facilitate fibrillation (*19, 20*). The presence of polymer coatings was confirmed by atomic force microscopy (AFM) (Figure S3) and X-ray photoelectron spectroscopy (XPS) (Figure S4 and S5).

Upon adsorption to the polymer coating, Fn spontaneously organized into a fibrillated network on PEA while remaining globular on PMA (Figure 1A, B). In order to ascribe any differences in adhesion to the conformation and not the amount of Fn, we analysed the quantity of protein on the two surfaces. XPS analysis determined that the amount of adsorbed protein was similar on the two polymer surfaces (Figure 1C and D, Table S1). The XPS survey scan and high-resolution C_1s_ XPS plots are shown in Figure S4 and Figure S5. The conformational differences of adsorbed Fn on the two coatings was corroborated by Fourier-transform infrared (FTIR) spectra, in which the peak positions indicate that Fn adsorbed to PMA adopts a mostly antiparallel β-sheet type secondary structure (*21*), similar to the globular, solution-state spectrum, while Fn on PEA adopts a more extended parallel β-sheet type structure (Figure 1E, Figure S6 and S7).

**Figure 1:**
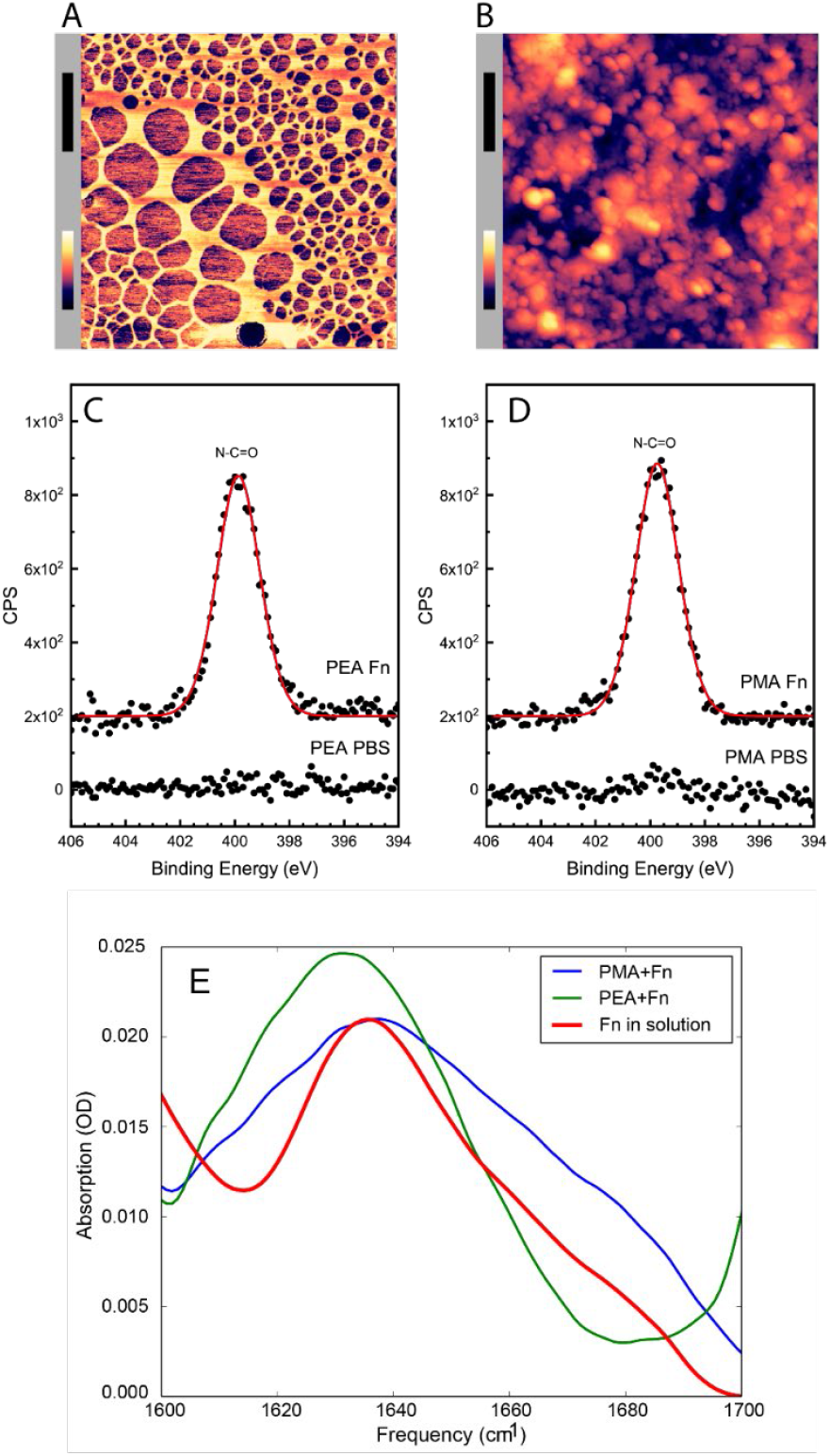
Adsorbed Fn remains globular on PMA and fibrillates on PEA-coated surfaces. AFM imaging shows the structure of adsorbed Fn on A) PEA and B) PMA (xy scale bar (blaock) = 500 nm, height scale bar (color) =115 nm). XPS analysis of the samples show similar chemical composition of Fn adsorbed to C) PEA and D) PMA, indicating that two polymer surfaces are both covered by Fn. E) FTIR spectral shape and intensity confirms that Fn adsorbed to PMA is similar to Fn in solution.

After validating the model system, Embp-mediated bacterial attachment to fibrillated and globular Fn was measured using a flow-cell system where the number of attached bacteria was counted by microscopy. Very few bacteria attached to the polymer coatings in the absence of Fn, and only fibrillated Fn stimulated attachment of *S. carnosus* expressing Embp_5F or Embp_9FG, (Figure 2A). Fn consists of two nearly identical subunits linked by a pair of disulfide bonds at the C terminal (*22*). Each subunit consists of three domains; F1, F2, and F3 (*23*). The globular and compact conformation of Fn is sustained by intramolecular electrostatic interactions between F1 1^st^-5^th^, F3 2^nd^-3^rd^, and F3 12^th^-14^th^ repeat (*24, 25*). Binding sites in these regions remain buried in the globular conformation; however, upon fibrillation on a surface or tissue interface, these binding sites become exposed (*26*). Since Embp only binds to fibrillated Fn, we hypothesize that it interacts with epitopes that are buried in the globular conformation, but become exposed when Fn fibrillates. Indeed, it was previously reported that *S. epidermidis* binds near the C terminal of Fn (*27*), and studies of recombinant Fn verified the interaction between Embp and the 12^th^ repeat of the F3 domain (*11*). This repeat may be one of several interaction points and has not been confirmed in full-length Fn adsorbed in its natural conformation. To test the interaction between the 12^th^ repeat of the F3 domain and the Fn-binding F- and FG-repeats, we repeated cell adhesion analysis on Fn-coated PEA after blocking the C-terminal heparin-binding domain II (F3 12^th^-14^th^ repeat) with antibody sc-18827. Control-samples were blocked with antibody F0916 specific for the F3 5^th^ repeat (Figure 2B). Blocking the F3 12^th^-14^th^ repeat decreased the adherence of *S. carnosus* by approximately 62 % for Embp_5F and 64 % for Embp_9FG (Figure 2B), supporting that Embp interacts with this subdomain. As the adherence was not completely abolished by blocking the Fn binding site, we cannot exclude the possibility that Embp interacts with other epitopes in Fn. However, the F3 12^th^-14^th^ repeat is of major significance.

**Figure 2:**
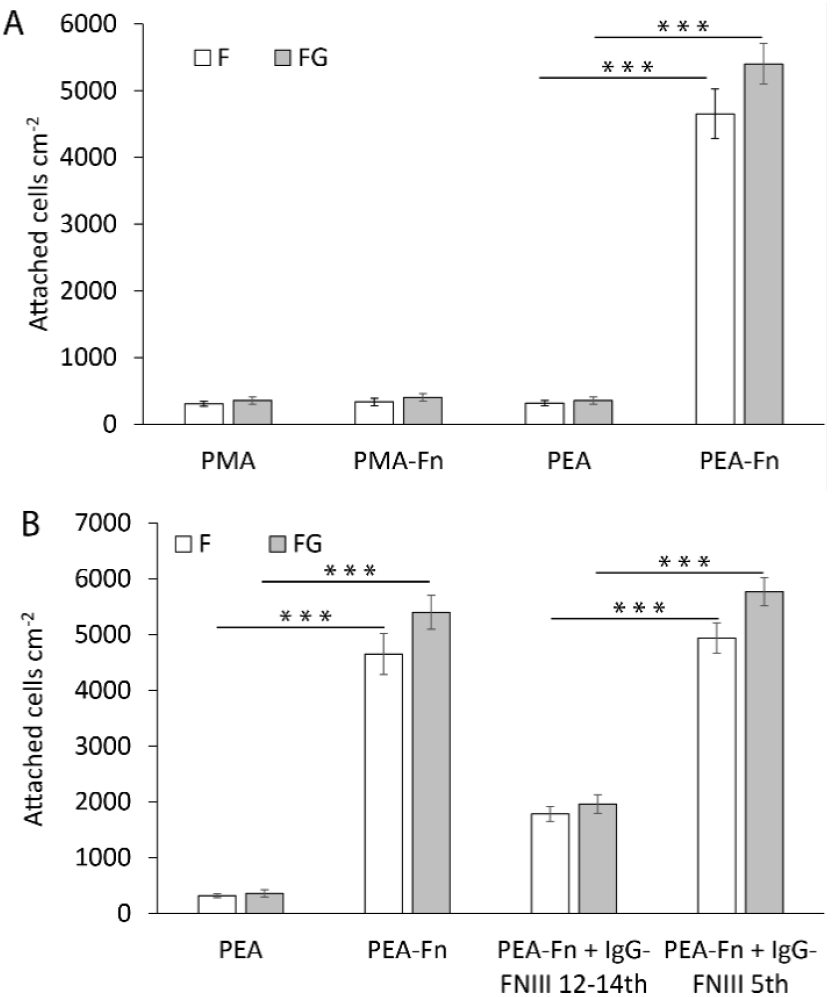
Embp only mediates bacterial attachment to fibrillated Fn. A) *S. carnosus* TM300 expressing either F- or FG-repeats were passed through flow cells for 2 hours before enumeration of attached cells by microscopy. Adsorbed Fn only promoted attachment on PEA-coated surfaces where Fn fibrillated. B) Identification of the Fn domain involved in bacterial attachment. The F3 12^th^-14^th^ repeats were blocked with a specific antibody prior to exposing bacteria to the surfaces. Blocking of the F3 5^th^ domain was included as a control for non-specific blocking of Fn by the antibodies. Values are averages from three independent experiments (error bars = S.D.). The two-tailed P-value from t test is < 0.0001.

### F and FG modules attach to fibronectin

After learning that Embp interacts exclusively with fibrillated Fn, we probed the strength of this interaction by single-cell atomic force spectroscopy. Single *S. carnosus* expressing Embp_5F or Embp_9FG were attached to colloidal AFM probes, approached to a Fn-coated PMA or PEA surface with controlled force, and then retracted to detect the force needed to detach the cell from the surface. As expected, the force-distance curves obtained from these experiments show that both F and FG fragments bind to fibrillated but not to globular Fn. The average maximum adhesion force between *S. carnosus* and surfaces with fibrillated Fn was 1.19 ± 0.21 and 1.16 ± 0.18 nN, respectively, for *S. carnosus* expressing Embp_5F or Embp_9FG (Figure 3). In contrast, the corresponding adhesion force to surfaces with globular Fn was only 0.16 ± 0.09 nN and 0.12 ± 0.04 nN. The adhesion force and the shape of the force-distance curves reflect multiple binding events between the cell and the Fn-coated surface. The multiple binding events could either be due to multiple Embp fragments on the cell surface interacting with Fn, or multiple interactions between a single Embp fragment and Fn.

**Figure 3:**
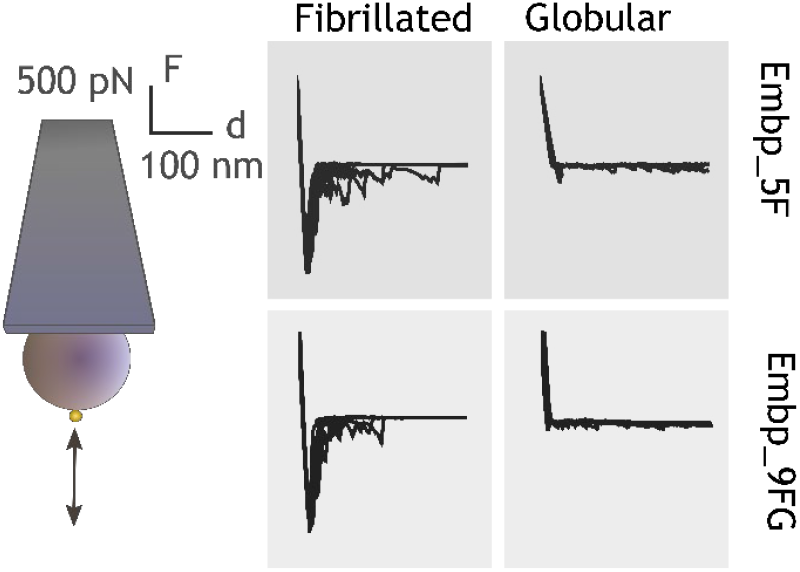
Single cell force spectroscopy shows that both the F- and FG-repeats adheres strongly to fibrillated Fn. Single *S. carnosus* cells expressing either F- or FG-repeats were immobilized on a colloidal AFM cantilever, and force-distance curves were measured by approaching and retracting the cantilever to surfaces with fibrillated or globular Fn. Adhesion events are recognized as negative peaks on the force axis below the horizontal baseline.

### Embp binds to fibrillated fibronectin in a Velcro-like manner

Embp contains 50 Fn-binding repeats, and it must be costly for *S. epidermidis* to produce this enormous 1 MDa protein. How might *S. epidermidis* benefit from the many repetitive binding-domains? We hypothesize that multivalent interactions can occur if the ligand for this giant adhesin is fibrillated, resulting in presentation of multiple binding domains in close proximity. Such multivalent binding would work like Velcro, as many weak binding events result in strong attachment. Such a Velcro-effect could provide adhesion forces strong enough to attach *S. epidermidis* to Fn via a single Embp protein. To investigate this hypothesis, we expressed and purified recombinant Embp fragments that contained 1, 4, and 15 repeats of FG-repeats, attached them to an AFM cantilever using 6His-NTA interaction, and quantified their interaction with Fn by single-molecule force spectroscopy. In agreement with previous experiments, the FG-repeat did not interact with the globular form of Fn (Figure 4). The interaction force of a 1 or 4 FG-repeat with fibrillated Fn was also insufficient to be detected. However, the interaction force of 15 FG-repeats was 432 ± 48 pN with fibrillated Fn, confirming the value of multi-domain interaction with the fibrillated ligand (Figure 4).

**Figure 4:**
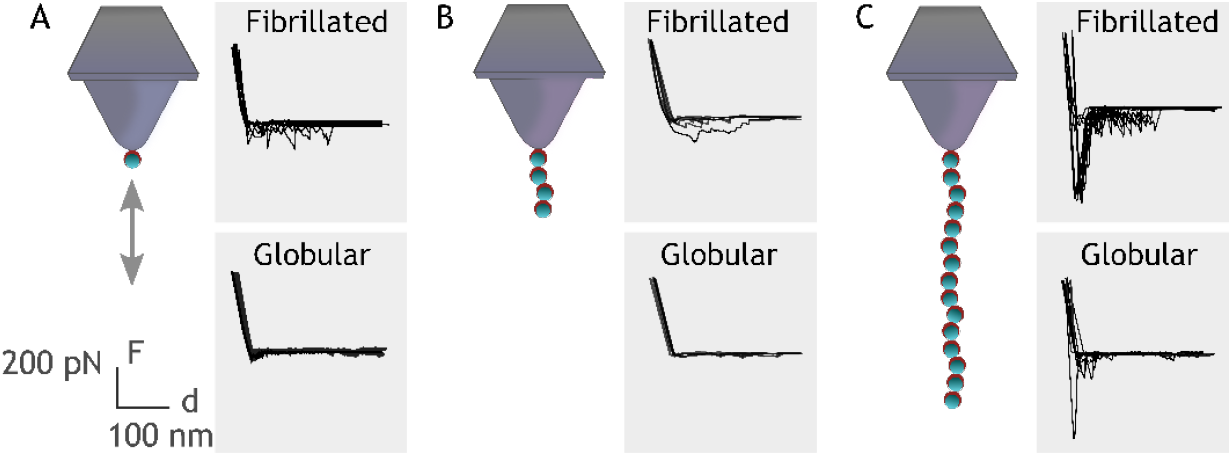
Single molecule force spectroscopy shows that multiple FG repeats are needed to detect binding to Fn. Recombinant Embp consisting of either 1 (A), 4 (B) or 15 (C) FG-repeats were tethered to a chemically modified silicon probe trough 6× His-NTA interaction. Force distance curves were measured towards fibrillated Fn on PEA and globular Fn on PMA.

### Embp is necessary for attachment under high flow

In our investigating of the interaction mechanism between Embp and Fn, we used fusion proteins that contained only a few of the F- or FG-repeats displayed on the surface of *S. carnosus* which has no other adhesive proteins. However, *S. epidermidis* has many other cell wall anchored adhesins, and the key to understanding Embp’s role in *S. epidermidis*’ pathogenicity therefore lies in understanding the circumstances under which Embp-producing *S. epidermidis* strains have an advantage. If oriented perpendicular to the cell surface, Embp could potentially stretch several hundred μm from the cell surface. We measured the hydrodynamic radius of *S. epidermidis* over-expressing Embp, and confirmed that it was significantly larger than for *S. epidermidis* lacking Embp (2.3 ±0.4 μm vs 1.3 ±0.2 μm, two-tailed t-test, n=3, P<0.001). We speculated that Embp would be more effective than other adhesins when *S. epidermidis* is attaching to Fn under high sheer stress. We therefore compared attachment of the two strains at low flow (1 mL min^-1^, 1.8 dyn cm^-2^) and high flow (18 mL min^-1^, 31.7 dyn cm^-2^), representative of the shear stress in arteries. At low flow, Embp did not affect attachment to Fn, but at high flow, attachment was only possible in the strain expressing Embp (Figure 5).

**Figure 5.**
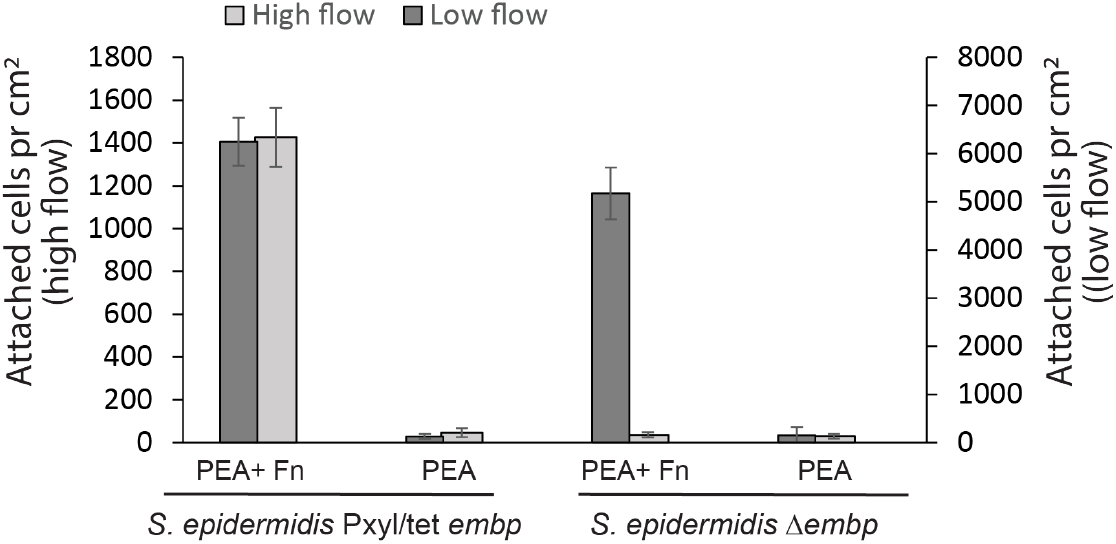
Embp is required for attachment at high flow. Attachment of *S. epidermidis* over-expressing Embp was compared to *S. epidermidis* lacking Embp at two different flow rates. Neither strain attached to PEA, and both strains attached to adsorbed Fn at low flow (dark grey bars). At high flow (light grey bars), only the Embp-expressing strain could attach to the adsorbed Fn.

## DISCUSSION

In this study, we show that the giant cell-surface protein Embp exclusively binds to fibrillated Fn because the binding site located at F3 12^th^ -14^th^ repeat is not accessible in the globular, soluble form of Fn. This discovery has implications for our understanding of how *S. epidermidis* colonizes host tissue and biomedical implants. Colonization and biofilm formation is the only virulence factor of *S. epidermidis*, and it is therefore imperative to cause disease (*28*). Implants provide a surface for attachment and are therefore vulnerable to biofilm infections. The implant surface is immediately covered by host proteins when it is inserted into the body, and bacterial attachment is assumed to occur via specific receptor-ligand interactions between adhesins on the bacterial cell and host proteins adsorbed to the implant surface. The abundance of the same host proteins in solution poses a dilemma: How can bacteria attach to implant surfaces via proteins that are also available in solution? Based on previous research (*10, 11*), we hypothesized that adsorption-induced conformational changes can affect how accessible a host protein is for bacterial adhesins. In the case of Embp and Fn, the fibrillation of Fn on the implant surface is decisive for its availability as a ligand for bacterial attachment. The selective interaction with fibrillated Fn illustrates how biology has solved the need for interaction with a protein in one location (the extracellular matrix of host tissue) while ignoring the same protein in another location (the bloodstream).

Our results also challenge notion that adsorbed host proteins always assist bacterial colonization. Motivated by this assumption, much research has been devoted to developing materials or coatings that prevent protein adsorption altogether. However, our results suggest that biomaterials’ susceptibility to biofilm do not only depend on protein adsorption, but also on the conformation of adsorbed proteins – a concept that is familiar to cell biologists studying adhesion of mammalian cells, but not to microbiologists studying bacterial attachment. The role of host protein conformation in bacterial attachment could explain the conflicting results from *in vitro* studies of how serum proteins affect attachment of staphylococci. Some studies report increased attachment (*11, 29-31*), while others report the opposite (*32, 33*). Perhaps these discrepancies reflect differences in how the underlying material affected the conformation of adsorbed serum proteins. Our study investigated bacterial attachment in a simplified model system with only one host protein, but if the concepts hold true in serum, it opens a door to controlling bacterial attachment by manipulating the conformation of adsorbed proteins, which is a much more manageable task than avoiding protein adsorption entirely.

The large number of repetitive domains in Embp and its homologs in *S. aureus* (Ebh) and *Streptococcus defectivus* (Emb) is unusual among bacterial adhesins, and we wondered how pathogens benefit from producing such large proteins. One benefit could be that the adhesin protrudes from the cell surface, which makes it easier to overcome electrostatic repulsion and reach its ligand. At present, no experimental evidence is available to directly demonstrate the organization of Embp on the cell surface. However, given the stretched overall architecture revealed by structural analysis (*10*), it appears plausible the molecule’s length is similar to that of Ebh in *S. aureus*. Ebh is a 1.1 MDa protein with 52 repetitive domains, and its length was predicted to be 320 nm (*34*). We show that the hydrodynamic radius of *S. epidermidis* over-expressing Embp was approximately 1 μm larger than *S. epidermidis* lacking Embp. It is thus likely that Embp extends well beyond the electric double layer.

Another advantage of producing adhesins with repetitive binding domains is the possibility for multivalent interaction with the ligand. Such multivalency is only possible if the ligand is fibrillated and thereby presenting many copies of its binding domain in close proximity of each other. Indeed, we showed that a single FG-module interacts weekly with Fn, while the interaction force of 15 FG-repeats was 432 ± 48 pN. We therefore propose that the selective interaction with fibrillated Fn is caused by i) the exposure of an otherwise buried binding domain in the fibrillated protein, and ii) the possibility of a stronger, multivalent interaction between multiple FG-repeats and the fibrillated ligand.

The typical strength of receptor-ligand bonds is about 20-200 pN (*35*). Hence, the interaction between 15 FG-repeats and Fn resulted in a very strong bond, and the interaction with full sized Embp is likely much stronger. The force required to detach Embp from fibrillated Fn will depend on whether all binding repeats detach at once, or whether detachment can occur from the serial unbinding of the repeats one by one. This is akin to detachment of Velcro: The attachment is very strong, but detachment can be obtained by the serial unbinding of individual interactions. The force-distance curves can provide information about how detachment occurs. On a close inspection, it is clear that initial retraction events are the strongest and provide the highest adhesion force, indicating that multiple bonds are broken at the same time (Figure 4B). But we also observe multiple subsequent perturbations as evidence to serial rupture, unfolding or stretching events. This holds true for interactions with 15 FG-repeats, but multiple perturbations are also observed for a single FG-repeat. In this case, the perturbations likely arise from the stretching and ruptures of surface bound fibronectin yielding an average adhesion force of 92 ± 27 pN, which is around the detection limit for this setup. Similar results were observed for 4 repeating units with average adhesion force of 68 ± 43 pN. On surfaces with globular Fn, little to no interaction was observed, although 15 repeating units undergo certain irregular interactions. An explanation for the observed irregularities could be simply the protein’s self-occupation/entanglement as would often happen with long polymer chains. Additionally, as can be seen from Figure 1, the surface distribution of Fn can vary, depending on the local density. This may be an insignificant problem for the dynamics of cell populations, however, when it comes to assessing individual cell’s or proteins’ behavior, the local density can affect the results.

In conclusion, the specific binding domain and repetitive structure of Embp provides an opportunity for *S. epidermidis* to interact selectively and strongly with fibrillated Fn, using a single adhesive protein. The open question is how this unique Velcro-like of interaction plays into the pathology of *S. epidermidis* and other pathogens that contain homologs of this protein in their genome. The genome of *S. epidermidis* is highly variable, and not all isolates possess the same repertoire of genes for host colonization and biofilm formation (*36*). Embp, however, is present in two thirds of *S. epidermidis* isolates from orthopedic device-related infections (*36*) and 90 % of isolates from blood stream infections (*9*), which indicates some importance for its pathogenicity. Strong attachment via a single protein could be particularly advantageous in locations where high shear forces make attachment difficult, such as in the blood stream. Biofilms generally do not form in blood vessels, unless there is an implant or a lesion on the endothelium. Infections like endocarditis often start with such a lesion (*37*), which makes the site susceptible to bacterial attachment. Fibrillated Fn forms on the surface of platelets in the early stages of wound healing (*38*), and perhaps the abundance of fibrillated Fn plays role in the elevated infection risk. We show that Embp is required for attachment of *S. epidermidis* under high flow (Figure 5), pointing to a role for Embp in attachment of *S. epidermidis* e.g. to cardiovascular grafts. Future research in animal models will determine Embp’s role in host and implant colonization in the cardiovascular system.

## MATERIALS AND METHODS

### Bacterial strains

*S. epidermidis* 1585 WT is a clinical isolate obtained from Rohde lab in UKE, Hamburg. *S. epidermidis* 1585Pxyl/tet *embp* (Embp overexpressed), *S. epidermidis* Δ*embp, S. carnosus* TM300 x pEmbp_5F (expressing 5 F-repeats) and *S. carnosus* TM300 x pEmbp_9FG (expressing 9 FG-repeats) were generated previously in the Rhode lab (*10*).

### Immunofluorescence of Embp fusion protein

Expression of Embp fusion protein in a non-adhesive surrogate host was critical for studying the interaction of Embp without interfering interactions from other adhesive proteins on the surface of *S. epidermidis*. We therefore started out by confirming the presence of Embp fragments on the surface of the surrogate host. *S. carnosus* TM300 WT, and *S. carnosus* TM300 x pEmbp_5F were grown overnight in brain heart infusion (BHI) broth with 10 μg ml^-1^ chloramphenicol (Sigma-Aldrich, Germany). Expression of Embp fragments in the mutant strains was induced with 200 ng ml ^-1^ anhydrotetracycline (AHT) after diluting the culture 100 times in BHI. Cells were grown for 6 hours at 37°C in a shaking incubator with 180 rpm until the 600 nm optical density (OD_600_) reached approximately 1. Cells were harvested by centrifugation (4000 × *g* or 10 minutes) and resuspended in phosphate buffered saline (PBS). A droplet of the resuspended cells was placed on a SuperFrost Ultra Plus slides (Invitrogen, USA) for 45 minutes to allow the bacteria to adsorb. After washing off unbound cells, the bacteria were fixed with 4% paraformaldehyde for 30 minutes at room temperature and washed twice with PBS. Samples were blocked with 5% goat serum (Invitrogen, USA) for 45 minutes, washed, incubated with anti-Embp2588 IgG antibodies (*39*) diluted 1:200 in blocking buffer at room temperature for 1 hour, washed three times, and then incubated with the secondary antibody (Anti-rabbit IgG conjugated with Alexa Fluor 635, Invitrogen, USA) diluted 1:300 in blocking buffer for 1 hour at room temperature. Cells were then washed three times and stained with 10 µM SYTO 9 (Invitrogen, USA) in PBS for 10 minutes, washed 3 times, and visualized by confocal laser scanning microscopy (CLSM) (LSM700, Zeiss, Germany) using 488 excitation for SYTO 9, and 639 nm excitation Alexa Fluor 635 conjugated antibody, and a 63x Plan-Apochromat N/A 1.4 objective.

### Interaction of Embp with soluble Fn

We first investigated if *S. epidermidis* or a surrogate host expressing Embp fragments interacted with soluble Fn in its globular conformation. *S. aureus* 29213 WT (positive control for soluble Fn binding) *S. epidermidis* 1585 WT, *S. epidermidis* 1585Δ*embp* (Embp knockout), were grown in BHI without antibiotics. *S. epidermidis* 1585Pxyl/tet *embp* (Embp overexpressed) was grown in BHI with 5 μg ml^- 1^ erythromycin, *S. carnosus* TM300 x pEmbp_5F (5 F-repeats) and *S. carnosus* TM300 x pEmbp_9FG (9 FG-repeats) were grown in BHI with 10 μg ml^-1^ chloramphenicol. Expression of Embp, F- and FG-repeats in the mutant strains was induced with 200 ng ml ^-1^ anhydrotetracycline (AHT) after diluting the culture 100 times in BHI. Cells were grown for 6 hours at 37°C in a shaking incubator with 180 rpm until reaching OD_600_ of approximately 1. Cells were harvested by centrifugation (4000 g for 10 minutes) and resuspended in PBS. A droplet of the resuspended cells was immobilized on a SuperFrost Ultra Plus slides (Invitrogen, USA) for 45 minutes to allow the bacteria to adsorb. The unabsorbed cells were removed by washing with PBS, and the adsorbed cells were then blocked with 3% BSA for 45 minutes. Cells were then incubated with 100 µg ml^-1^ Fn in PBS (Sigma-Aldrich, F0895) for 60 minutes at room temperature. The unbound Fn was removed by washing three times with PBS. The samples were then fixed with 4% paraformaldehyde for 30 minutes at room temperature. Immunolabeling was then performed as described above, Anti-Fn mouse IgG (Sigma-Aldrich) diluted 1:100 in blocking buffer and the secondary antibody (Anti-mouse IgG conjugated with Alexa 635, Goat IgG - Invitrogen) diluted 1:300 in blocking buffer. Cells were stained and prepared for imaging as described above.

### Preparation of polymer-coated surfaces

Quantification of interaction forces between Embp and Fn in its globular or fibrillated form would require that Fn was immobilized to a surface. We used a previously published model system (*20, 40*) to generate Fn-coated surfaces that displayed Fn in these two conformations, while the physicochemical properties of the underlying surface was very similar, namely PEA and PMA. Polymers of ethyl acrylate and methyl acrylate were synthesized from their monomers (99% pure, Sigma-Aldrich, Germany) using radical polymerization. Benzoin (98% pure, Sigma-Aldrich, Germany) was used as a photoinitiator with 1 wt % for PEA and 0.35 wt % for PMA. The polymerization reaction was allowed in Schlenk flasks exposing to ultraviolet light (portable UV lamp with light of 390-410nm) up to the limited conversion of monomers (2 hours). Polymers were then dried to constant weight in a vacuum oven at 60 °C for 12 hours. Both polymers were solubilized in toluene (99.8% pure, Sigma-Aldrich) to concentration of 6% w/v for PEA and 2.5% w/v for PMA. 2 hours sonication in an ultrasonic bath at room temperature was used to make the polymer soluble. Glass slides (76 × 26 mm, Hounisen) were cleaned with sonication in ultrasonic bath for 15 minutes in Acetone, Ethanol, and Milli-Q water respectively, and then dried under nitrogen flow. A thin film of polymer solution was coated on clean slides using spin-coater (Laurell Technologies) with acceleration and velocity of 1000 rpm for 30 seconds. The spin-coated films were degassed in a desiccator for 30 minutes under vacuum and then put in a vacuum oven at 60 °C for 2 hours to remove toluene.

### Fn adsorption to PMA and PEA

A hydrophobic marker (PAP pen – Sigma-Aldrich) was used to draw a small circle (around 0.5 cm square area) on the spin-coated slides. Fn from human plasma (Sigma-Aldrich, F0895) was dissolved in PBS at concentration of 20 µg ml^-1^ and 100 μl sample was adsorbed on each slide for 1 hour at room temperature.

### Atomic force microscopy for imaging of Fn adsorbed to PMA and PEA

Experiments were conducted on three replicate samples with JPK Nanowizard IV (JPK, Germany) using HQ: CSC38/No Al (Mikromasch, USA) and TR400PSA (Asylum Research, USA) cantilevers. We used the fluid mode of operation to visualize Fn adsorbed to PEA and PMA without introducing artifacts from sample drying. The operating environment was controlled in a closed liquid chamber at 21°C with minimal evaporation. The operation parameters were set to optimize resolution with minimum possible damage or artifact from contaminations on the tip. Typically, scans were started with a large scan area of minimum 10× 10 µm^2^ with a rather low scan resolution of 64 × 64 pixels and a high scan rate > 1 Hz. Once an area of interest was identified, a higher resolution image 256 × 256 or 512 × 512 pixels of a smaller scan are was acquired at lower scan speeds (< 1 Hz). The acquired data was processed using Gwyddion open software (http://gwyddion.net/) for necessary corrections of tilt etc.

### XPS of Fn adsorbed to PEA and PMA

A100 μl of Fn (20 µg ml^-1^) was adsorbed on a polymer spin-coated 1 × 1 cm glass slides (these slides were cut manually in the chemistry lab workshop) for 1 hour, and samples were then washed three times with Milli-Q water and dried under N_2_ flow. The chemical composition of the adsorbed layer was analyzed with a Kratos AXIS Ultra DLD instrument equipped with a monochromatic Al K*α*X-ray source (hν = 1486.6 eV). All spectra were collected in electrostatic mode at a take-off angle of 55°(angle between the sample surface plane and the axis of the analyzer lens). The spectra were collected at new spots on the sample (n=3, 1 replicate) and were charge corrected to the C_1s_ aliphatic carbon binding energy at 285.0 eV, and a linear background was subtracted for all peak areas quantifications. Analyzer pass energy of 160 eV was used for compositional survey scans of C_1s_, O_1s_, N_1s_, Na_1s_, Si_2p_, Cl_2p,_ P_2p_, and K_2p_. High-resolution scans of C_1s_ and N_1s_ elements were collected at an analyzer pass energy of 20 eV. Compositions and fits of the high-resolution scans were produced in CasaXPS. The data is presented in a table as an average and standard deviation of the three sample spots.

### FTIR analysis of Fn adsorbed to PEA and PMA

FTIR measurements were performed on a Bruker Vertex v70 with 128 scans per spectrum and a 7 mm diameter beam spot. The concentration of Fn (20 µgml^-1^) results in very small IR absorbances of the polymer layers, so therefore, the spectra of stacks of 8 coated CaF2 windows were measured simultaneously. For this, 8 spin coated CaF2 window surfaces with PMA and PEA were incubated for one hour with the 20 µgml^-1^ Fn solution in PBS prepared in D_2_O (d-PBS hereafter), after rinsing the surfaces with d-PBS and placed 4 sets of windows (spaced by 25 µm Teflon spacers that were filled by d-PBS, with the polymer and protein-coated sides submerged in the d-PBS) in a custom-made IR cell. The incubated sample spectra were background-corrected by subtracting the spectra of the same neat d-PBS loaded windows. Before subtraction, we (i) corrected for small differences in the overall transmission of the protein and background samples (due to e.g. small differences in the amount of scattering of the IR beam, which can become significant with 8 consecutive windows) by subtracting the absorption at 7500 cm^-1^, and (ii) corrected for small differences in the exact water-layer thickness by scaling the spectra using a spectrally isolated absorption band of the D2O (the v1 + v2 combination band of the solvent’s OD-bending and stretching mode1 at 3840 cm^-1^) to determine the scaling factor. But there is no reason to assume that the water and polymer layers thicknesses are related. Therefore, the resulting background-corrected amide-I (1600-1700 cm^-1^) PEA+Fn spectrum (Figure 1) still contains a tail of the 1733 cm^-1^ ester peak, which is absent in the resulting PMA+Fn spectrum. This is (i) because there is approximately 6 times more PEA present than PMA (as indicated by a least-square fit that minimized the total intensity of the subtraction of the PMA from the PEA background spectra in the 1700-1760 cm^-1^ region, see supplementary materials Figure S6, S7 and the accompanying supplementary materials text), and (ii) because the Fn incubation results in a slight loss of polymer, which is impossible to compensate for well by subtraction of the polymer spectra, because the 1733 cm^-1^ ester peak shape is affected by the presence of the protein (see figure S6(d)). The broadening of this peak by Fn incubation is probably because the ester groups in contact with the protein are slightly shifted with respect of the more buried ester groups that are not changed by the protein adsorption, resulting in two subpeaks that are slightly offset in frequency. Even though the PMA layer appears to be thinner and/or less dense, it will probably still be composed of many monolayers (as indicated by the XPS measurements), so this difference in thickness is not expected to affect the protein’s interfacial behavior.

### Quantification of bacterial attachment under flow

Ibidi sticky-slide VI 0.4 chambers (Ibidi, Germany) were glued to polymer-coated glass by using an equivalent mixture of silicon (DOWSIL 732 - Dow corning) and UV activating glue (Loctite 3106 Light Cure Adhesive). After flow cell assembly, 50 μl of Fn (20 µg ml^-1^) dissolved in PBS was injected to the channel of a flow cell and allow to adsorb statically for 1 hour at room temperature. The unbound Fn was removed by a flow of PBS (6 ml hour^-1^) using a syringe pump (Harvard Apparatus, USA) for 15 minutes. Bacterial cells were subcultured from an overnight culture and grown for 6 hours at 37°C and 180 rpm, harvested by centrifugation (4000 g for 10 minutes), and resuspended in PBS to an OD_600_ of 0.1. The cell suspension was flowed through the flow cell chamber at 3 ml hour^-1^ for 2 hours at room temperature. The unbound cells were washed with PBS at 9 ml hour^-1^ for 30 minutes. Attached bacteria were visualized by brightfield microscopy (Zeiss Axiovert A100, 20x objective) and counted. A minimum of 5 images were acquired per replicate, and a minimum of 500 cells were counted per replicate.

The first experiment compared attachment of *S. carnosus* TM300 x pEmbp_5F and *S. carnosus* TM300 x pEmbp_9FG to PEA and PMA surfaces with and without Fn to investigate if the Fn-binding domains of Embp interacted selectively with the fibrillated form of Fn. The second experiment investigated which domain of Fn Embo interacted with. Previous studies had shown that that Embp binds to the F3 12^th^-14^th^ domain of Fn (*11*), however, this experiment was only performed with recombinant fragments of Fn and not the full length protein. We therefore investigated the role of this Fn domain in the attachment of bacteria via Embp. Flow-cell experiments were carried out as described above, comparing attachment via F og FG to fibrillated Fn directly or after blocking for F3 12^th^-14^th^ domain withn IgG antibodies (Anti-Fn, sc-18827, Santa Cruz biotech). As a control, fibrillated Fn was blocked with IgG antibodies specific for another Fn domain (F3 5^th^ domain). The unbound antibodies were removed with PBS (6 ml hour^-1^, 15 minutes) before investigating bacterial attachment as described above.

The final experiment addressing attachment under flow compared the attachment of *S. epidermidis* 1585 Pxyl/tet *embp* and *S. epidermidis* Δ*embp*. The strains were inoculated from single colonies into BHI broth (amended with 5 μg ml^-1^ erythromycin and 200 ng ml^-1^ anhydrotetracycline (ATC) for the Pxyl/tet *embp* strain) and grown overnight at 37°C, 180 rpm, harvested by centrifugation, and resuspended in PBS to OD_600_ = 0.3, transferred to the syringe pump and passed through the flow-cells at either 1 mL min^-1^ or 18 ml min^-1^ flowrate for 1 h followed by a 30 min PBS washing step of 6 ml min^-1^ or 36 ml min^-1^, respectively. Attached cells were imaged by brightfield microscopy and by CLSM after staining with 20x SYBR Green II (Sigma Aldrich).

### Hydrodynamic radius

*S. epidermidis* 1585 Pxyl/tet *embp* and *S. epidermidis* Δ*embp* were prepared as described above, transferred to cuvettes and analysed by dynamic light scattering (DLS) (Folded Capillary Zeta Cell, malvern US). Measurements of surface charge and cell diameter was carried out using Zetasizer Nano (Malvern Panalytical).

### Single-cell force spectroscopy

Single-cell force spectroscopy (SCFS) measurements were conducted on Fn adsorbed in its globular conformation to PMA or its fibrillated conformation to PEA. For SCFS measurement, colloidal probes with 10 µm glass beads (SHOCON-BSG-B-5, Applied NanoStructures Inc., USA) were selected and coated by polymerizing a dopamine solution of 4 mg ml^-1^ dopamine hydrochloride (99%, Sigma-Aldrich, H8502) in 10 mM Tris-HCl buffer at pH 8.5, and then calibrated *in situ* for single-cell attachment. *S. carnosus* TM300 expressing 5 F- and 9 FG-repeats were subcultured from overnight cultures and incubated for 6 hours in fresh media, harvested and resuspended in PBS as described above. A 100 µl drop of this solution was placed on a glass slide and incubated for 10 minutes, after which the unadsorbed bacteria were removed by rinsing with PBS. A colloidal probe was immersed and positioned on top of a single cell with the help of inverted optical microscope. The probe was made to contact a single cell for 5 minutes then retracted after the cell attachment. Once a cell was picked up (confirmed by optical microscopy), the substrate was changed to Fn coated surfaces and SCFS was executed. The acquired force-distance plots were processed using the Nanowizard’s (JPK, Germany) own processing software. Experiments were conducted on two replicate samples.

### Cloning and purification of F- and FG-repeats

Genomic DNA was extracted from *S. epidermidis* 1585 WT strain using the Qiagen DNA kit (Qiagen, Hilden, Germany) by following the instructions of the manufacturer. The only exception made in the kit protocol was that cells were lysed with 15 U of lysostaphin, which was added to buffer P1. The nucleotide sequence of one, four, and fifteen repeats of the FG-repeats were amplified from genomic DNA using primers (Table S2) with Phusion High-Fidelity PCR Kit (NEB - E0553S). The PCR products were purified with GenElute PCR Clean-Up Kit (Sigma-Aldrich NA1020). The expression vector pET302/NT-His was digested with EcoR1 restriction enzyme (NEB R0101S) and run on a 1.5 % agarose gel. The digested vector was purified from the gel using the GenElute Gel Extraction Kit (Sigma-Aldrich NA1111). The purified PCR product of each recombinant Embp (rEmbp) was ligated with the digested vector in a ratio of 3:1 using the Gibson assembly ligation matrix mix (NEB E5510S). Each ligation reaction was incubated for 1 hour at 50 °C. The ligated products were transformed into the chemically competent *E. coli* strain (Top10). The colony PCR was performed with REDTaq ReadyMix (Sigma-Aldrich R2523) using the T7 promoter primer as forward and the T7 terminator primer as the reverse. Cells from each selected colony were grown overnight in LB with 100 μg ml^-1^ ampicillin, and a plasmid miniprep was prepared using GeneJET Plasmid Miniprep Kit (Thermo Scientific K0702). Plasmids were sequenced with both T7 promoter and terminator primers by the Eurofins A/S (Hamburg, Germany). The plasmid of each Embp construct was transformed into an expression system (chemically competent *E. coli*, BL21-DE3). A single colony of the transformants was used to inoculate 2 L of LB with 100 μg ml^-1^ ampicillin until the OD_600_ of 0.6. For the overexpression, cells were induced with 1M IPTG, and incubated for 16 hours on 28 °C in shaking incubator at 180 rpm. Cells were harvested and lysed in binding buffer with sonication (30% amplitude, 15 seconds off, 15 seconds on) for 3 minutes on ice. After centrifugation, the supernatant was filtered with a 0.22 μm syringe filter and run on a Nickel-Nitrilotriacetic Acid (Ni-NTA) column using ÄKTA Purifier-10 purification system. The column was washed with 5 - 8 column volumes, and the fusion proteins Embp was then eluted in fractions using elution buffer. Fractions of each rEmbp were pooled, concentrated with Amicon Ultra centrifugal filter tubes with a cut-off 3 kDa (Millipore Sigma UFC9003). Proteins were further purified with (HiTrap Q FF) column by anion exchange (IEX) chromatography using IEX binding and elution buffer, followed by size exclusion chromatography (SEC) with column (Superdex 200 Increase 10/300 GL) using MES buffer on ÄKTA Purifier-10 purification system. After each column, the elution fractions were run on SDS-PAGE to check the purification quality. Buffers used for rEmbp purification are listed in Table S3.

### Single-molecule force spectroscopy

Single-molecule force spectroscopy (SMFS) measurements, similar to SCFS, conducted on Fn adsorbed to either PEA and PMA. For the SMFS experiments, the probes were prepared by attaching rEmbp fragments of various lengths with the use of His_6_-NTA interaction. The procedure was similar to that of Obataya *et al (41)*. In short, silicon probes were cleaned with ozone, and UV light then kept in one to tone isopropyl alcohol and ethanol mixture overnight. Tips were rinsed in deionized water and air-dried, after which they were functionalized with 2% (3-mercaptopropyl) trimethoxysilane in EtOH for 30 minutes. Probes were then exposed to Maleimide-C3-NTA in 50% DMF/100 mM Tris-HCl (pH7.5 ± 0.1) overnight. 10 mM NiCl_2_ was used to chelate the NTA groups on the tip, which was then dipped in bovine serum albumin (1 mg ml^-1^ in PBS) to passivate the surface. The attachment of 6X His-tagged rEmbp fragments with 1, 4 or 15 repeating units of Fn binding repeats was completed by 1 hour incubation of respective samples at room temperature. After a probe for each repeating unit were prepared, force-distance curves were collected on three replicate samples and processed, as mentioned above.

## Supporting information

Supplementary results

## ACKNOWLEDGEMENTS

This work was funded by the Carlsberg Foundation, Grant number CF16-0342. Cecilie Siem Bach-Nielsen is gratefully acknowledged for quantification of *S. epidermidis* attachment under high and low flow rates. TWG and SJR thank the Lundbeck Foundation for postdoc fellowships.

## Notes

### Competing Interest Statement

The authors have declared no competing interest.

### Summary of Updates

The Title, Introduction, Results and Discussion sections have been revised for greater clarity.

