## Supplementary results for "The giant staphylococcal protein Embp facilitates colonization of surfaces through Velcro-like attachment to fibrillated fibronectin"

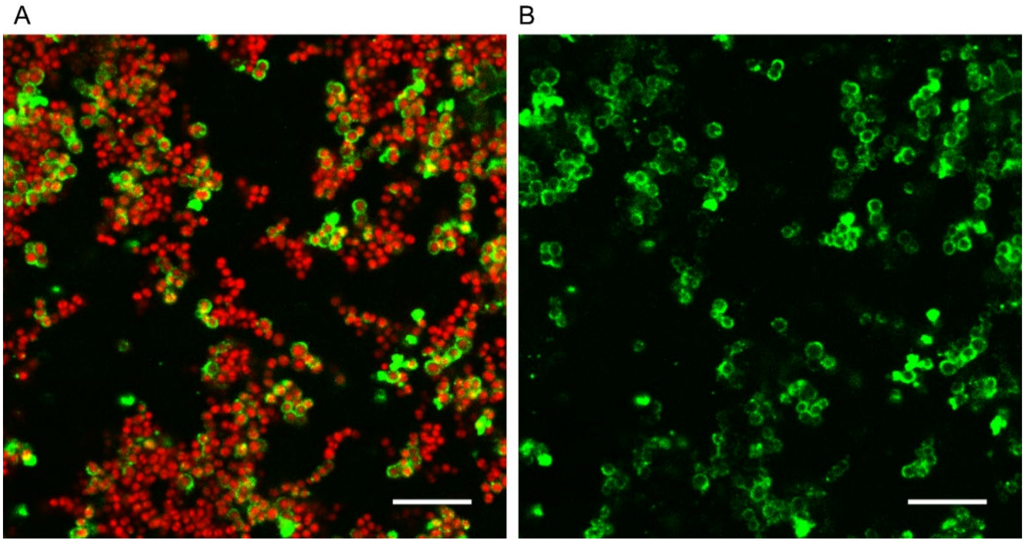


**Figure S1:**
Localization of recombinant Embp (5F repeats) on the surface of *Staphylococcus carnosus* TM300. Bacteria were stained with SYTO 9 (red), and recombinant Embp was labeled with anti-Embp antibodies followed by anti-Rabbit secondary IgG conjugated with Alexa Fluor 635 (green). The scale bar is 10 µm. A) Overlay of bacteria (red) and Embp (green), B) Embp only (green)

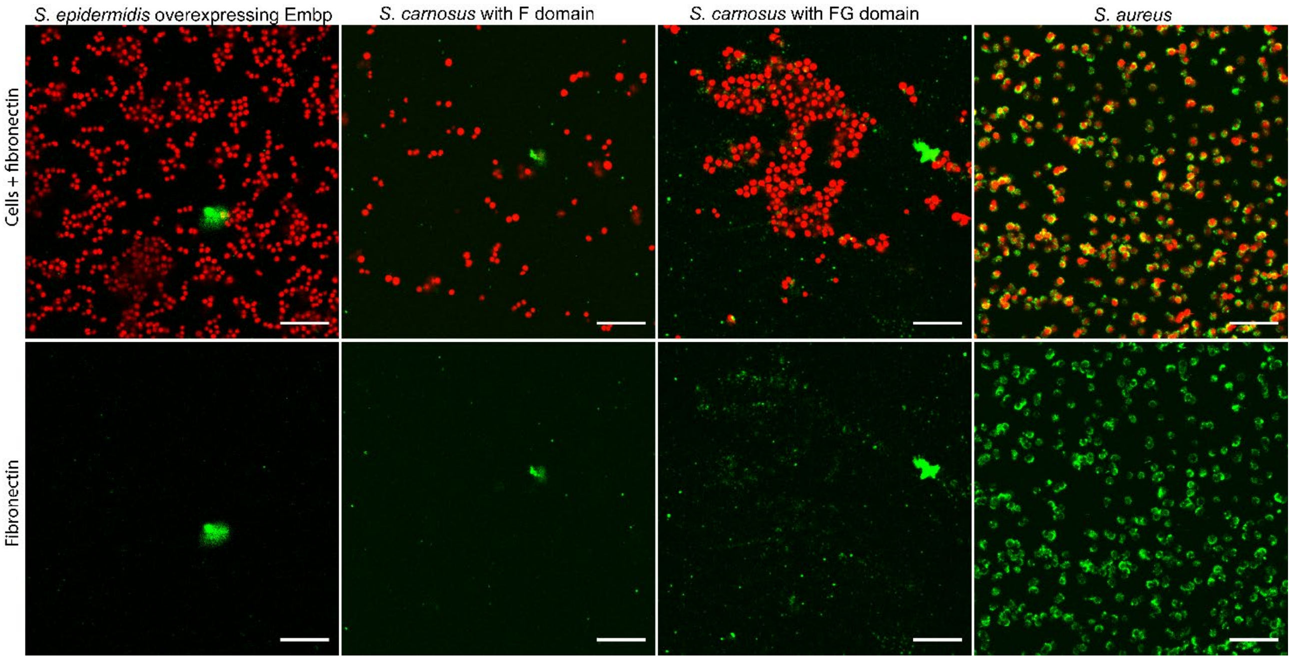


**Figure S2:** Interaction of soluble Fn with *S. epidermidis* 1585Pxyl/tet *embp* overexpressing Embp or *S. carnosus* TM300 expressing recombinant Embp (5 F or 9 FG repeats). *S. aureus* was used as positive control. Bacteria were stained with SYTO 9 (red), and recombinant Embp was labeled with anti-Embp antibodies followed by anti-Rabbit secondary IgG conjugated with Alexa Fluor 635 (green). Top panel shows overlay of bacteria (red) and Fn (green). Bottom panel shows Fn only (green). Scale bar = 10 µm.

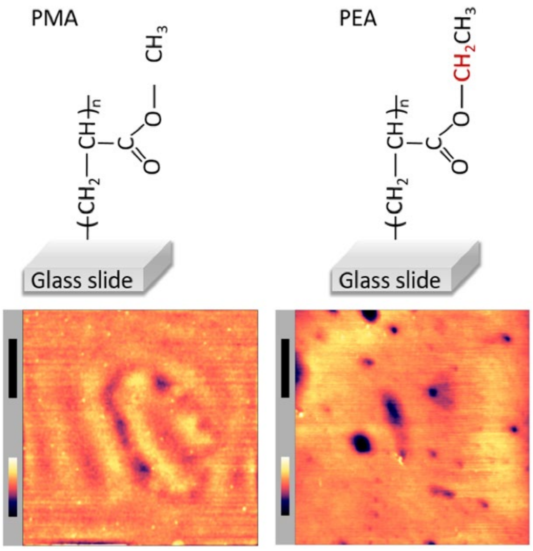


**Figure S3:** Structure of poly (methyl acrylate) (PMA), and poly (ethyl acrylate) (PEA) surfaces. AFM images (bottom panel) of PMA- and PEA- coated glass surfaces were acquired in PBS. The xy-scale bar (black) is 1 µm and z-scale bar (in color) is 26 nm for PMA and 16 nm for PMA.

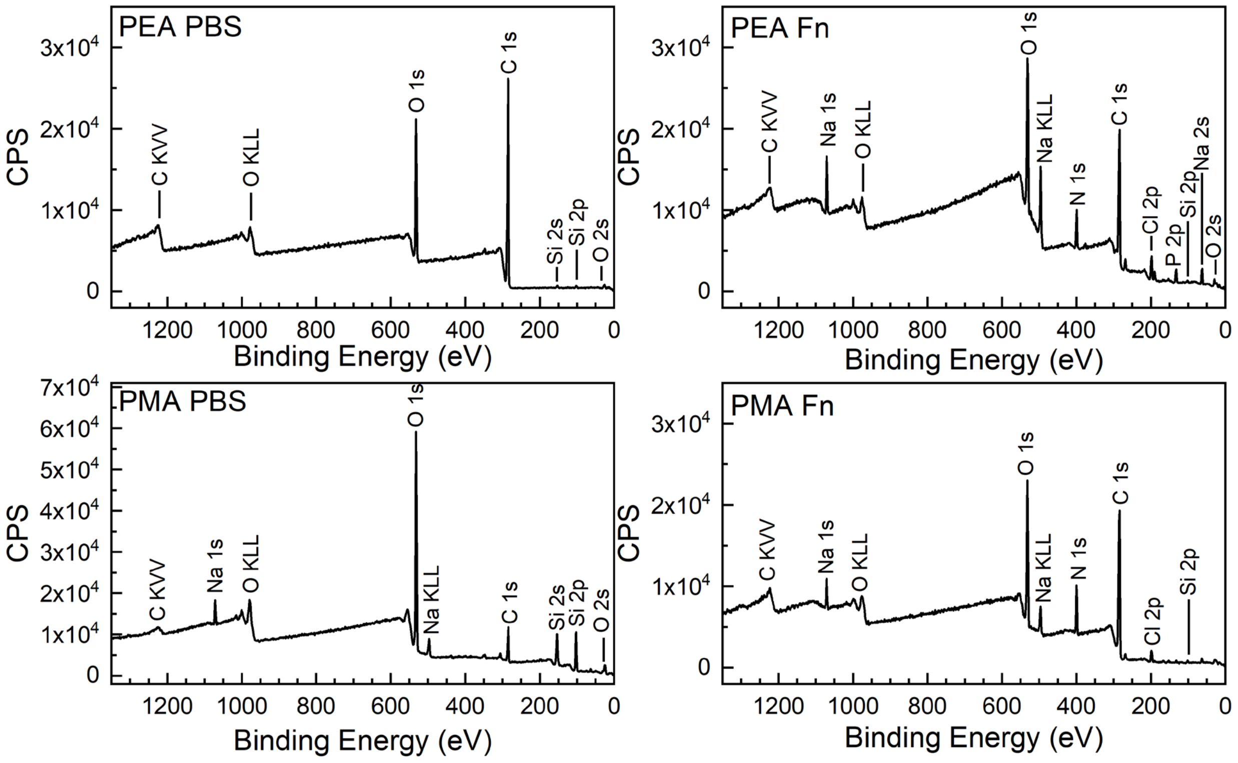


**Figure S4:** Plots of XPS survey spectra of PEA and PMA polymer with either PBS added or fibronectin. The survey scans allow for the determination of the elements present at the sample surface.

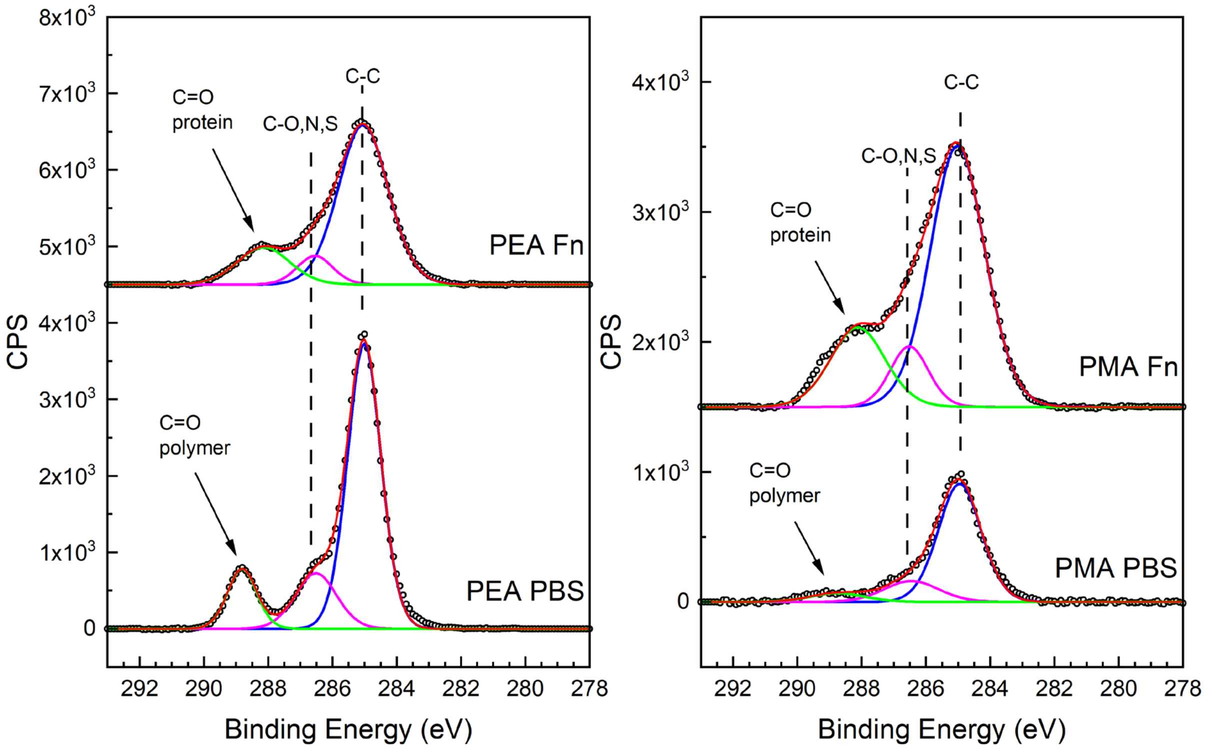


**Figure S5:** High-resolution $C_{1s}$ XPS plots of peaks characteristic of protein addition on polymer surfaces. (bottom) Spectra of only PEA or PMA polymer incubated with PBS. (top) Spectra of PEA and PMA polymer incubated with PBS and fibronectin (Fn). The C1s high-resolution spectrum have peaks at 285.0 eV, 286.5 eV, 288.0 eV, and 289.0 eV assigned to the C-C, C-O, N, or S, the C=O in a protein, and the C=O in a polymer, respectively.

### Additional information about the FTIR experiments

Infrared spectroscopy of the amide-I region (1600-1700 $cm^{-1}$) is a secondary-structure sensitive technique[2, 3]. The coupling between the amide-I modes of backbone amide groups is strong, due to the large transition-dipole moment of this mode, leading to unique spectral signal signatures of most different types of secondary structures. Here, we applied the technique to study samples consisting of stacks of 8 layers of PEA or PMA after incubation with a 20 μg/mL Fn solution for one hour (see main text Figure 1, and the main text Materials and Methods section for the details of the background correction). Here, additional information about the FTIR experiments is provided, as well as additional insights that can be obtained from these spectra by analyzing them in various ways (see below).

Under the excitonic approximation[4, 5], the area of the amide-I region spectra is conserved for different secondary structures, if the concentration remains the same. Therefore, the similarity in the area of the Fn-incubated PMA and PEA spectra corroborates (within the experimental error of the rinsing step and the spectral background correction) the XPS data that indicated that the amount of bound protein is similar for both polymer surfaces. The amide-I area of the Fn-incubated PEA samples is only 7% smaller than the Fn-incubated PMA.

Furthermore, the peak frequencies of the spectra indicate that the PMA-incubated Fn adopts a mostly antiparallel β-sheet type secondary structure, while the PEA-incubated Fn samples appears to adopt a more extended parallel β-sheet type structure[2]. The high-frequency (~1685 $cm^{-1}$) absorption, indicative of antiparallel β-sheets[2], that is present in the PMA-incubated samples is also present in the solution-phase spectrum, and the low-frequency (~1636 $cm^{-1}$) absorption maxima of PMA-incubated and solution-phase Fn have similar frequencies. Hence, PMA-incubated Fn probably adopts a similar conformation as in solution. The remaining spectral differences suggest that there are more random-coil and turn structures[3] (1640-1675 $cm^{-1}$) in the PMA-incubated Fn. In contrast, the PEA-incubated Fn lacks the high-frequency absorption, which indicates parallel β-sheet structure[2], and the red-shifted low-frequency mode (now at ~1630 $cm^{-1}$) indicates that the β-sheets are delocalized over more β-strands, which could be due to the formation of intermolecularly hydrogen-bonded structures[6, 7].

*Additional insights from various processed versions of the FTIR spectra*

In supplementary Figure S6, the raw spectra and various processed versions are depicted. The raw spectra, depicted in Figure S6(a), show that the scattering is different for all 4 samples, because the spectra are offset with respect to each other. This is expected for the current sample, with the 8 consecutive windows and the fact that the polymer layers scatter the IR light in a not fully reproducible manner. To demonstrate that the presence of protein can already be seen with minimal data processing, in Figure S6(b), the spectra are vertically shifted so as to obtain 0 OD at 1700 $cm^{-1}$ to show that also the sample thicknesses are not exactly the same, as can be seen by the small intensity difference of the combination of $\nu_2$ + the libration mode of $D_2O$ at 1555 $cm^{-1}$. Now also the minute amount of protein absorption in the amide-I region can be appreciated (see inset), with more high-frequency absorption in the PMA case, and more low-frequency absorption in the PEA case, although care must be taken with drawing direct conclusions from this as one first has to correct for the sample thicknesses (as one can see from intensities of the 1555 $cm^{-1}$ mode). Looking at the 1733 $cm^{-1}$ polymer ester mode, one can now see that the incubation with the Fn solution, and the rinsing with d-PBS removed a small amount of polymer, reducing the polymer thickness. When overlapping the post-incubation spectra to the pre-incubation spectra on the ester peak intensity (Figure S6(c)), it becomes apparent that in the PEA case 12% of the polymer layer is lost, and in the

PMA case only 3%. Finally, matching the intensity of the ester peak in all spectra to the peak intensity in the pre-incubation PEA spectrum indicates a factor ~7 between the amount of PEA and the PMA present at the samples (Figure S6(d)).

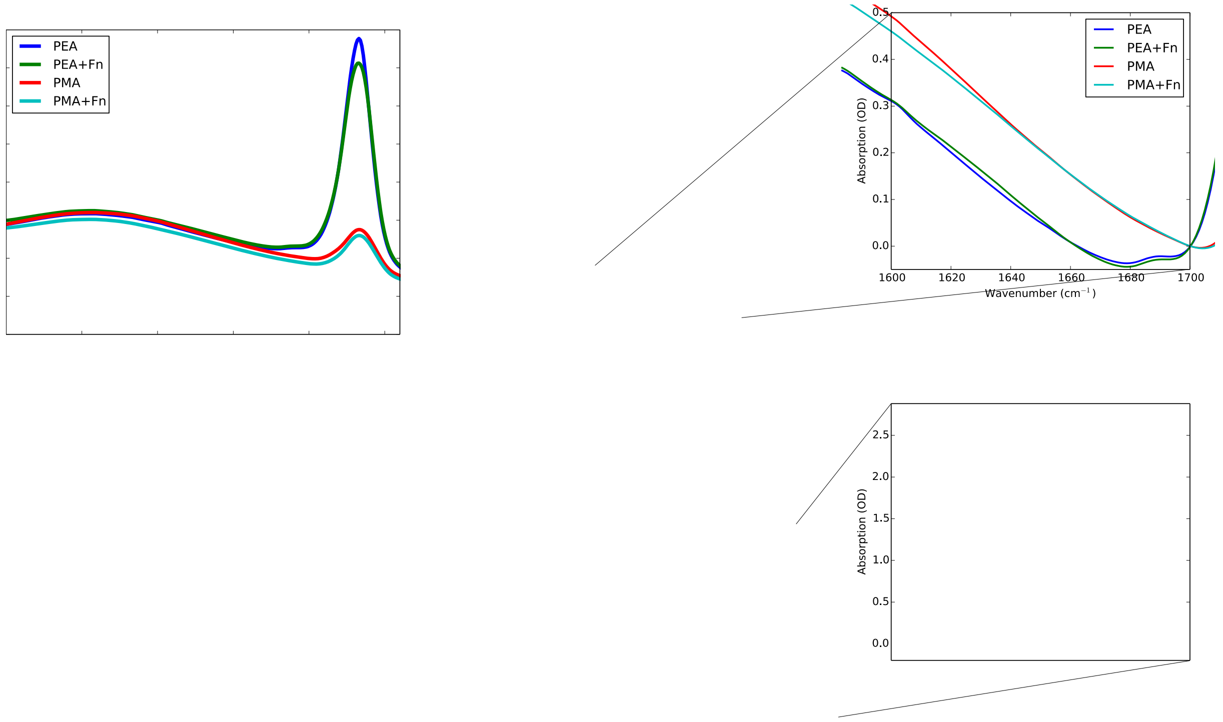


**Figure S6:** Various variants of processing of the FTIR spectra, indicating the relative contributions of scattering, protein absorption and polymer absorption (see associated supporting materials text).

A more accurate way to determine the relative amount of polymer-ester absorption, optimizing the overlap between the polymer ester peaks the pre- and post-incubation samples using a least-square fit with the offset and scaling factor as the fit parameters (instead of matching the intensities at 1700 and 1730 $cm^{-1}$ as done in Figure S6(d)). In supplementary Figure S7, the result of such a least-square fit of the polymer ester peak is given before (a) and after incubation (b) with 20 μg/mL Fn. In this procedure, performed with a home-written Python script, the difference between two spectra is minimized with a constant offset $c$ and a scaling factor $m$, i.e.: spectrum1(ν) – ($m$ * spectrum2(ν) + $c$). The constant offset will reflect differences in the scattering, while $m$ reflects the factor between the number of IR oscillators (when looking at the same, isolated) peak. When performing this procedure only in the 1700-1760 $cm^{-1}$ region both before and after the incubations, and subtracting the PMA spectra from the PEA spectra, we find factors of 5.3 and 6.4, respectively, i.e., there are approximately 6 times as many ester groups present in the PEA case, as compared to the PMA case.

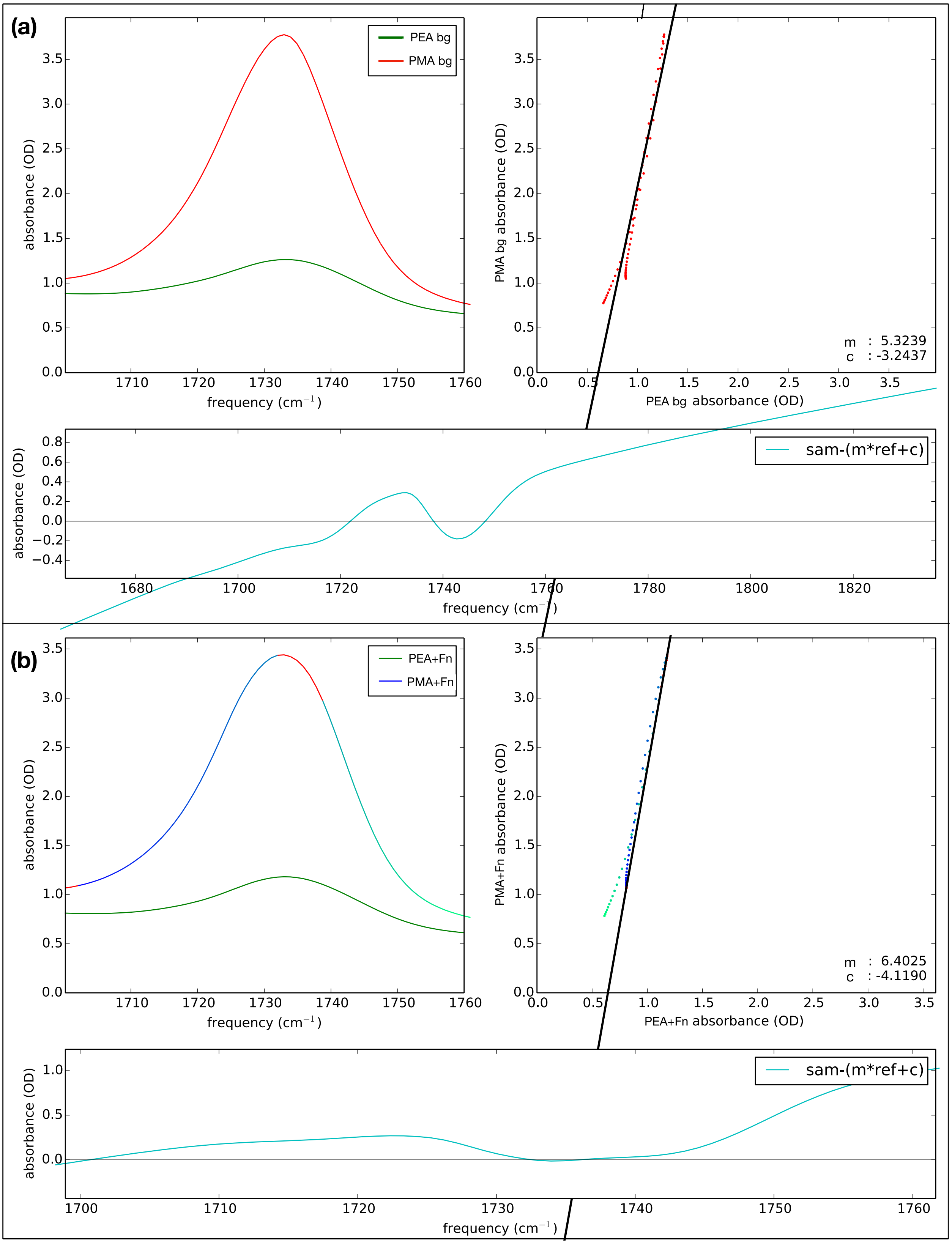


**Figure S7:** Results of least-squares fit before (top) and after (bottom) Fn incubation of the spectra of the PEA- to the PMA-coated surfaces, in order to obtain the relative number of PEA and PMA molecules on the surfaces (see associated supporting materials text).

**Table S1:** XPS survey spectrum atomic percent compositions for polymer PEA and PMA and either PBS or PBS with Fn

| Element | PEA PBS [%] | PEA Fn [%] | PMA PBS [%] | PMA Fn [%] |
|---|---|---|---|---|
| **C 1s** | 79.5 (1.0) | 55.8 (1.0) | 21.4 (1.4) | 67.4 (3.8) |
| **O 1s** | 19.6 (1.0) | 27.0 (1.1) | 55.5 (1.2) | 21.4 (1.3) |
| **N 1s** | n.d. | 8.3 (0.4) | n.d. | 7.7 (1.5) |
| **Si 2p** | 0.9 (0.4) | 0.1 (0.1) | 19.8 (0.8) | 0.2 (0.2) |
| **Na 1s** | n.d. | 3.6 (0.1) | 3.2 (0.7) | 1.9 (0.6) |
| **Cl 2p** | n.d. | 2.5 (0.3) | 0.1 (0.1) | 1.4 (0.3) |
| **P 2p** | n.d. | 2.7 (0.1) | n.d. | n.d. |
| **K 2p** | n.d. | n.d. | n.d. | n.d. |

Note: not detectable (n.d.); (.) = standard deviation; [%] = atomic percent

**Table S2:** Primers for rEmbp cloning into pET302/NT-His

| Primer name | Sequence 5´ to 3´ |
|---|---|
| FIVAR-GA forward | AGAAGGAGATATACATATGCATCATCATCATCATCACGTGGAATTCGAAAACC TGTATTTTCAGGGCGGAGATCAAAAACTTCAAGATG |
| 1 FIVAR-GA reverse | TCCGATTATACCTAGGCTCGAATATCATCGATCTCGAGCGGAATTCTTAATGA AGATTTTGTTCAGC |
| 4 FIVAR-GA reverse | TCCGATTATACCTAGGCTCGAATATCATCGATCTCGAGCGGAATTCTTAATGT AAACTTTCTCTAGC |
| 15 FIVAR-GA reverse | TCCGATTATACCTAGGCTCGAATATCATCGATCTCGAGCGGAATTCTTAATT TAACGATGTTTCTGC |
| T7 Promotor | TAATACGACTCACTATAGGG |
| T7 Terminator | GCTAGTTATTGCTCAGCGG |

**Table S3:** Buffer used for rEmbp purification

| Buffer name | Composition |
|---|---|
| Binding/lysis | 50 mM $K_2PO_4$, 500 mM NaCl, 400 mM imidazole, pH 7.4 |
| Ni-NTA Elution | 50 mM $K_2PO_4$, 500 mM NaCl, 40 mM imidazole, pH 7.4 |
| IEX binding | 20 mM BIS-TRIS propane, pH 6.0 |

| | |
|---|---|
| IEX elution | 20 mM BIS-TRIS propane, 1M NaCl, pH 6.0 |
| SEC | 50 mM MES, 150 mM NaCl, pH 6.0 |
